# Genomic evidence of Great Hornbill (*Buceros bicornis*) consuming the endemic Southern Hill Myna (*Gracula indica*) and first mitochondrial genome of *G. indica*

**DOI:** 10.64898/2026.11.23.768500

**Authors:** Pooja Yashwant Pawar, Rohit Naniwadekar, Jahnavi Joshi

## Abstract

Great Hornbill (*Buceros bicornis*) consumes animal prey, including bird chicks, but prey species are not easily identifiable from field observations to species level. Using genomic data recovered from a Great Hornbill faecal sample, we report consumption of an endemic Southern Hill Myna (*Gracula indica*), providing the first genomic evidence of this prey-predator interaction. The phylogenetic analyses ascertained the identification of *G. indica*. Additionally, we present the first annotated mitogenome of *G. indica*, providing a valuable genomic resource and confirming its phylogenetic placement within the family Sturnidae.

## Introduction

Asian hornbills are predominantly frugivorous birds. In addition to fruits, hornbills consume a wide range of invertebrate and vertebrate prey. Animals provide an important source of protein, especially during the breeding season. Studies on Asian hornbill diets during the breeding season have documented consumption of a wide range of animal prey, including invertebrates, amphibians, reptiles, birds, and mammals (Kannan & James, 1997; Mudappa 2000; Bora et al., 2023).

Great Hornbill (*B. bicornis*) is one of the largest Asian hornbills with the widest geographic distribution in the Western Ghats, Himalayan foothills, Indo-Burma region, and Sundaland to Sumatra (Kinnaird & O’Brien, 2008). Extensive breeding biology and foraging behavioural studies from India (Pawar et al., 2018) and Thailand (Ouithavon et al., 2005; Chaisuriyanun et al., 2011) demonstrate that a small but significant proportion of the Great Hornbill diet consists of animal prey. However, observational studies are limited in their ability to identify prey to species level. Consequently, vertebrate prey is usually identified only to broad taxonomic groups (e.g., birds, reptiles, or mammals), leaving species-level predator-prey interactions largely unknown (Ouithavon et al., 2005; Chaisuriyanun et al., 2011; Pawar et al., 2018; Bora et al., 2023). Non-invasive DNA obtained from fecal matter provides an opportunity to identify prey at the species level (Rodgers & Janečka, 2012).

Using mtDNA genomic data recovered from a Great Hornbill faecal sample, we confirm in this study the consumption of Southern Hill Myna *G. indica* by Great Hornbill. Additionally, we provide the first annotated mitochondrial genome of the endemic *G. indica* and examine its evolutionary relationship with closely-related species of the family Sturnidae. We also quantify mitochondrial sequence divergence between *G. indica* and its sister species *G. religiosa*, providing baseline molecular data for future phylogenetic and population genetic studies.

## Methods

### Sample collection and genomic data generation

A fresh fecal sample of Great Hornbill was collected beneath an active nest tree in Nelliampathy, Kerala, India, in February 2024. Following the direct observation of the nesting female defecating, a fecal sample was collected. It was preserved in Longmire’s lysis buffer and stored at room temperature during fieldwork. The sample was subsequently stored at -20□ until DNA extraction. DNA was extracted using Qiagen’s QIAmp Fast DNA Stool Mini Kit (Catalogue ID: 51604) following established protocols. DNA yield and quality were assessed using high-sensitivity dsDNA Qubit Kit on Qubit 4 Fluorometer (Thermo Fisher Scientific, USA, Catalogue no. Q33238), NanoDrop spectrophotometer (Thermo Scientific), and 4150 TapeStation system (Agilent Technologies, Inc. 2021) with D1000 DNA ScreenTape kit. The sequencing library was prepared using the TruSeq DNA PCR-Free kit (Illumina) following the manufacturer’s protocol at the NGS Facility in CSIR-CCMB, Hyderabad. The short-read (150 bp paired-end) sequences were generated on the Illumina NovaSeq 6000 sequencer to obtain whole-genome sequence data. The reads obtained from the sequencer were demultiplexed using bcl2fastq and quality filtered using Fastp v 0.23.4 (Chen, 2023). Reads having low Phred scores (below Q20) were removed.

### Mitogenome assembly and annotation

The trimmed raw reads were used to extract mitochondrial DNA reads using ‘--clade Chordata’ and ‘--genetic_code 2’ and assembled using the default assembler MEGAHIT in MitoZ v 3.6 (Meng et al., 2019). Using the same software, mitochondrial genome assembly was annotated and visualized. The taxonomic identity was assessed by comparing the assembled mitochondrial genome sequence assembled in this study using BLASTn search in the GenBank and on barcoding gene Cytochrome c oxidase subunit I (COI) against the BOLD database (http://www.boldsystems.org/; Ratnasingham et al., 2007).

### Intra-specific and inter-specific evolutionary divergence estimation

We downloaded published sequences for two genes (Cytochrome b and NADH dehydrogenase subunit 2) for *G. religiosa*, a sister species to *G. indica* (Miller et al., 2025), from NCBI’s GenBank repository (See Table 2). We used Cytochrome b (n = 5; sequence length = 1143 bp) and NADH dehydrogenase subunit 2 (n = 7; sequence length = 1040 bp) (Table 2). We estimated evolutionary divergence between *G. indica* and *G. religiosa* by calculating pairwise genetic distance using the Kimura 2 parameter (K2P) model, which accounts for differences in the rates of transitions and transversions using software MEGA7 (Kumar et al., 2016).

### Phylomitogenomic analyses

We aligned 13 protein-coding genes individually using the MUSCLE algorithm implemented in MEGA7 and subsequently concatenated the alignments. We reconstructed phylogenetic relationships among members of Sturnidae using maximum-likelihood (ML) and Bayesian inference (BI) approaches to ascertain the phylogenetic position of our sample. We downloaded mitochondrial genomes of species belonging to the family Sturnidae (mynas and starlings), as well as representatives of the closely related families Muscicapidae (flycatchers), Mimidae (thrashers, mockingbirds, and tremblers), and Buphagidae (oxpeckers), following Ray et al. (2023) and used as outgroups in the analyses (see Table 1).

**Table 1.** List of species and their GenBank accession numbers which were used for Phylomitogenomic analysis.

| **Species name** | **GenBank accession number** |
|---|---|
| *Gracula indica* | This study |
| *Acridotheres cristatellus* | JF810423 |
| *Buphagus erythrorhynchus* | NC_053077 |
| *Gracula religiosa* | JF937590 |
| *Leucopsar rothschildi* | NC_053050 |
| *Luscinia cyanura* | KF997864 |
| *Niltava davidi* | NC_039538 |
| *Phoenicurus auroreus* | KF997863 |
| *Rhabdornis inornatus* | NC_051020 |
| *Sturnus cineraceus* | HQ896037 |
| *Sturnus nigricollis* | JQ003191 |
| *Sturnus sericeus* | HM859900 |
| *Sturnus tristis* | NC_015195 |
| *Sturnus vulgaris* | KT946691 |
| *Toxostoma redivivum* | NC_053053 |

**Table 2.** List of GenBank accession numbers of *G. religiosa* for Cytochrome b and NADH dehydrogenase subunit 2 genes used in which were used in inter-specific evolutionary divergence estimation.

| **GenBank accession number** | **Location** | **Cytochrome b** | **NADH dehydrogenase subunit 2** |
|---|---|---|---|
| NC015898 | China | x | x |
| JF937590 | China | x | x |
| MN991547 | Museum specimen, USA | x | x |
| KJ456290 | Museum specimen, USA | x | x |
| DQ466862 | Captivity, USA | - | x |
| EF468158 | Captivity, Singapore | - | x |
| EF468159 | Captivity, Germany | - | x |
| KJ455447 | Museum specimen, USA | - | x |

We performed Maximum likelihood (ML) analyses on the concatenated dataset using ModelFinder implemented in IQ-TREE 2 (Nguyen et al., 2015). The best-fitting models were TIM+F+I+G4 for ATP6, TN+F+I for ATP8, TPM2u+F+I+G4 for COX1, TPM2u+F+G4 for COX2 and COX3, TVM+F+I+G4 for CYTB, GTR+F+G4 for ND1, TN+F+I+G4 for ND2, TIM+F+I for ND3, K3Pu+F+G4 for ND4L, TIM+F+I+G4 for ND4, TIM+F+I+R2 for ND5, and HKY+F+G4 for ND6. The branch support was assessed using the ultrafast bootstrap method with 1000 replicates.

We used PartitionFinder2 (Lanfear et al. 2016) to assess the best partition scheme and nucleotide substitution models from a set of only MrBayes models using the AICc criterion. The optimal partitioning scheme consisted of five partitions. GTR+I+G was selected as the best-fitting model for partitions 1–4, comprising (1) ATP6, ATP8, ND3, ND4L, ND1, ND2; (2) COI, COII, COIII; (3) Cytb; and (4) ND4, ND5, respectively. HKY+I+G was selected for partition 5 (ND6). This partitioning scheme and the corresponding substitution models were used for Bayesian analysis in MrBayes (Ronquist et al., 2009) where four chains were run for 50 million generations, with trees sampled every 5,000 generations. Model parameters, including nucleotide state frequencies, substitution-rate matrices, gamma-distributed rate variation, and the proportion of invariant sites, were unlinked among partitions, allowing each partition to have independent parameter estimates. Branch lengths were also unlinked among partitions, and a variable-rate prior was used to model rate variation among partitions. Convergence and mixing of the chains were assessed using Tracer v1.7.2 (Rambaut et al., 2018), with effective sample size (ESS) values >200 considered indicative of adequate sampling. We discarded the first 25% of sampled trees as burn-in and generated a consensus tree from the remaining trees. Phylogenetic trees were visualized using FigTree (Rambaut, 2009).

## Results and discussion

We obtained 10 GB of paired-end short-read data consisting of 72 million reads for the *B. bicornis* faecal sample and used 70 million reads after quality filtering. The mitochondrial genome was assembled from 11,150 mapped reads. The assembled mitogenome was 16,815 bp in length and exhibited 100% coverage with a mean sequencing depth of 95.05×. It consisted of 13 protein coding genes, two ribosomal RNAs and 22 tRNAs (Fig 1). The overall GC content is 47.57%.

**Figure 1.**
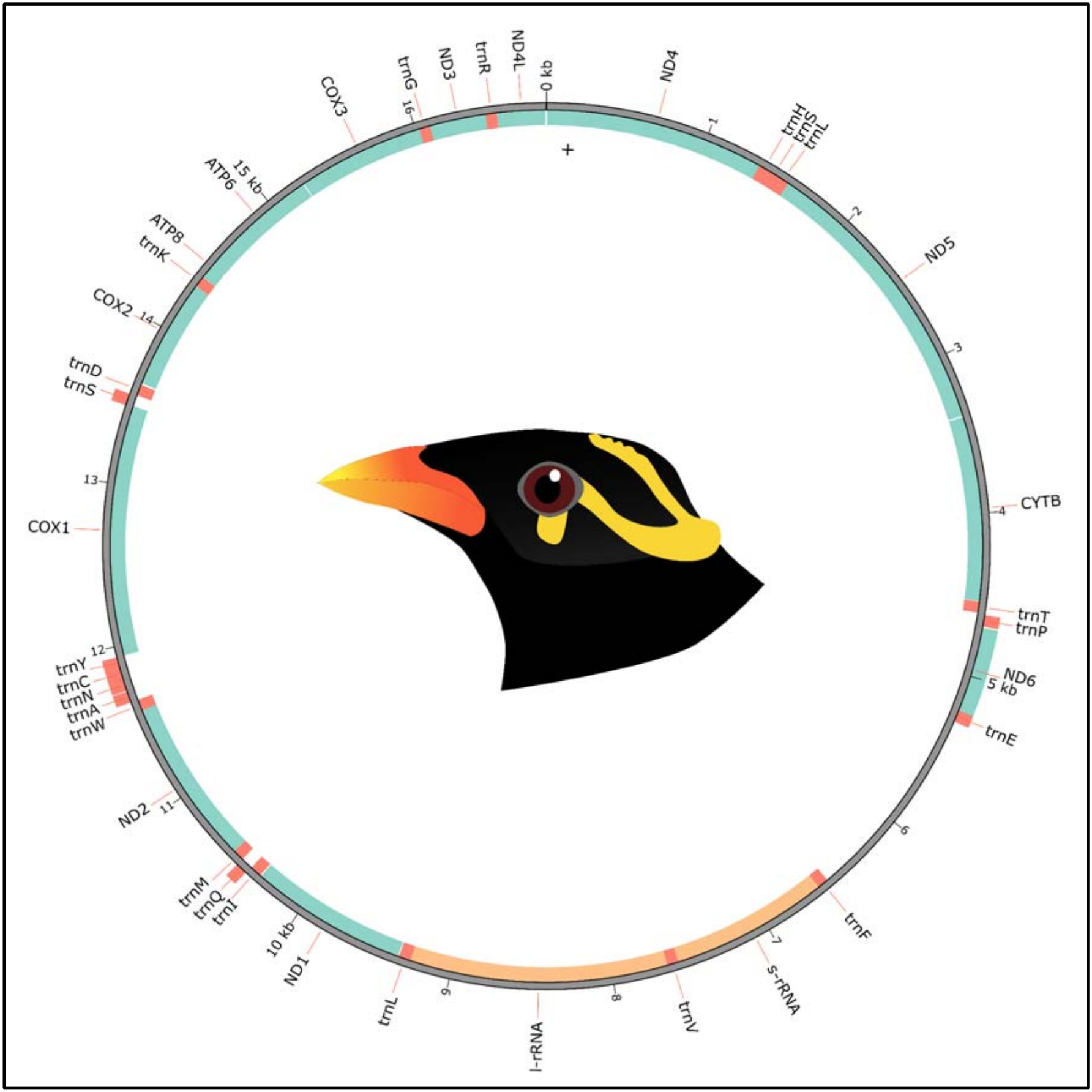
Annotated mitochondrial genome plot for *G. indica*. The genetic fragments in the plot indicate positions of protein-coding genes (green), rRNA (blue), and tRNA (orange). Illustration of *Gracula indica* by Shyamal, via Wikimedia Commons, used under CC BY-SA 3.0 license.

The assembled mitochondrial genome (16,815 bp) was queried against the NCBI nucleotide database using BLASTn. The sequence showed the highest similarity to *Gracula religiosa* (97.73% nucleotide identity, 100% query coverage, E-value = 0). Similarly, a search against the animal COI database in BOLD showed 96–97% similarity to *G. religiosa*. However, considering the known non-overlapping geographic distributions of these species, *G. religiosa* (distributed in Eastern Ghats, Himalaya and Southeast Asia) and *G. indica* (endemic to Western Ghats) (Craig and Feare, 2020; Limparungpatthanakij, 2022), we tentatively considered the recovered mitogenome to represent *G. indica*. Interspecific sequence divergence for cytochrome *b* (*Cytb*) and NADH dehydrogenase subunit 2 (*ND2*) ranged from 2.3% to 3.7%, whereas intraspecific divergence ranged from 0% to 0.3% (Fig. 3). The discrepancy between the molecular database matches and the expected species identity also highlighted the lack of *G. indica* in publicly available mitochondrial and COI reference databases.

Phylogenetic tree reconstruction based on 13 mitochondrial protein-coding genes, including 15 species of Sturnidae and representatives of Muscicapidae, Mimidae, and Buphagidae as outgroups, placed the genus *Gracula* in a distinct clade comprising *G. religiosa* and our putative *G. indica* (Fig. 2). The ML and BI analyses yielded identical topologies, with strong bootstrap support (100%) and posterior probabilities of 1 for the *Gracula* clade. Our sample was placed within the *Gracula* clade as sister to *G. religiosa*, supporting its placement within the genus.

**Figure 2.**
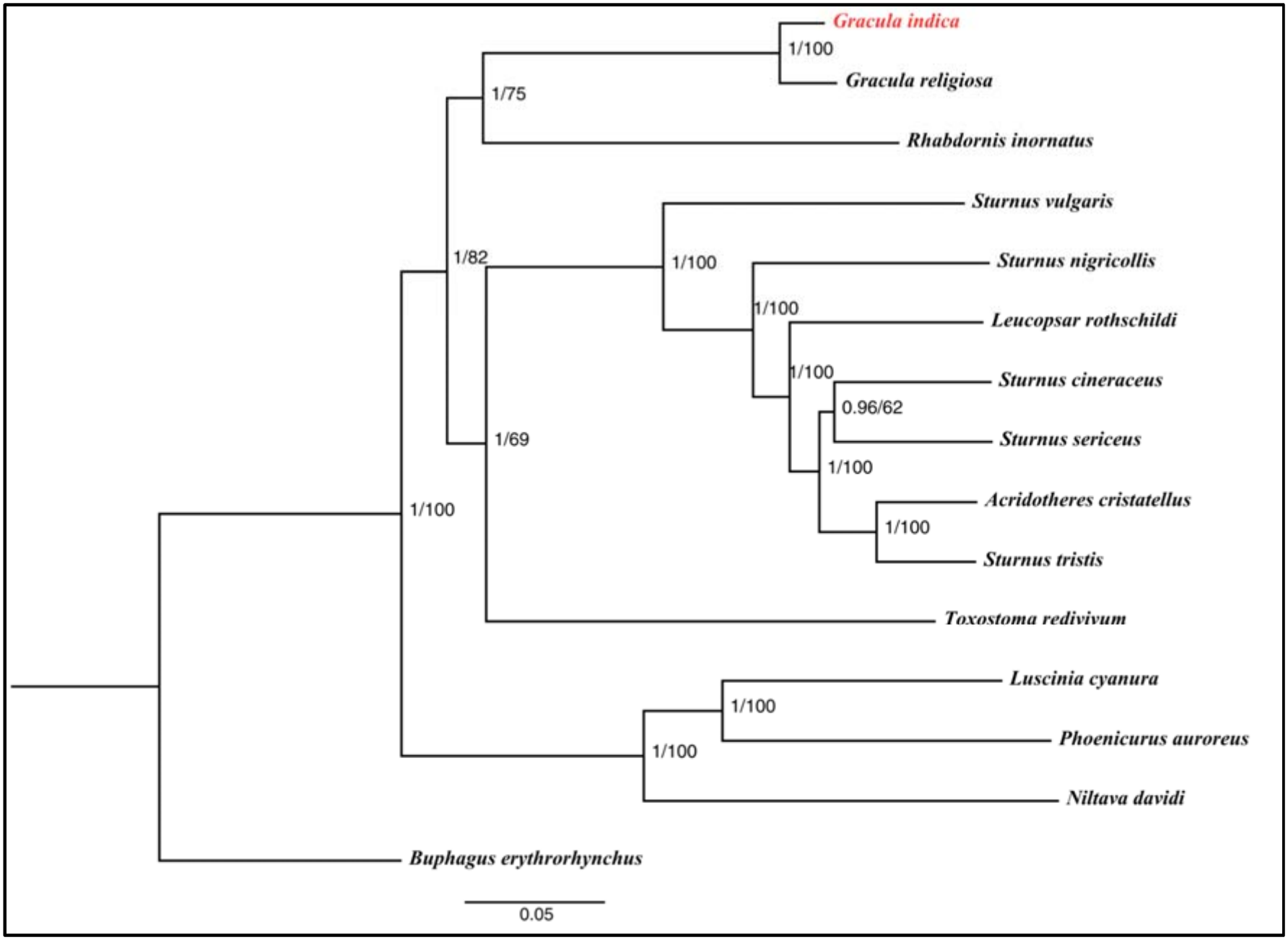
Phylogenetic relationship between *G. indica* (this study, in red) and members of the Sturnidae family. The node labels indicate posterior probability and maximum likelihood percentage, respectively.

**Figure 3.**
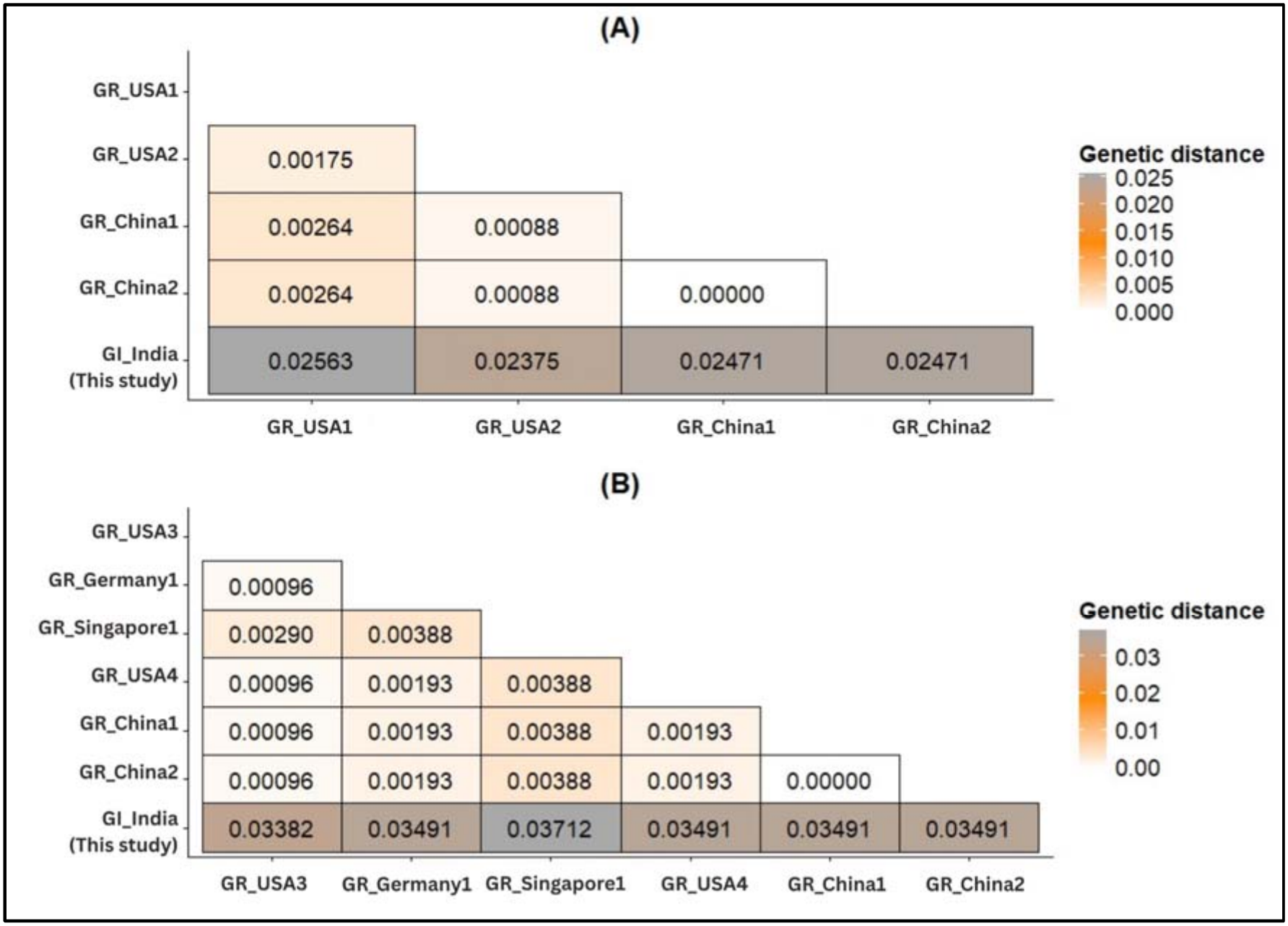
Evolutionary divergence between sister species of genus *Gracula*: genetic distance (p-distance) between individuals of *G. religiosa* (GR) and *G. indica* for (GI); (A) Cytochrome b gene (n = 5; sequence length = 1143 bp) and (B) NADH dehydrogenase subunit 2 gene (n = 7; sequence length = 1040 bp).

Taken together, the geographic location of the collected sample, its phylogenetic placement, and the observed genetic divergence suggest our sample could be of *G. indica*.

To our knowledge, this represents the first quantitative estimate of mitochondrial sequence divergence between *G. indica* and *G. religiosa*.

This provides the first genomic evidence that Great Hornbill consuming Southern Hill Myna *G. indica*, an endemic species in the Western Ghats. Direct observations on feeding behaviour of Great Hornbill had reported them to be predating on bird chicks (Ouithavon et al., 2005; Chaisuriyanun et al., 2011; Pawar et al., 2018), however it is extremely difficult to identify the prey species in the field. Hence, non-invasive sources of DNA such as feces provide an opportunity for robust identification of the prey species (Rodgers & Janečka, 2012).

The whole genome sequencing allowed us to further build on the existing molecular knowledge of the endemic *G. indica*. We provide a high-quality, complete, annotated mitochondrial genome of *G. indica (100% genome, average sequencing depth 95x)*. The overall GC content is 47.57%, similar to other species in the family Sturnidae (Qian et al., 2013; Ray et al., 2023). This study demonstrates that whole-genome sequencing of hornbill faecal samples can simultaneously identify prey species and generate genomic resources for poorly studied endemic taxa. The annotated mitogenome provides a genomic resource for future phylogenetic and population genetic studies of the endemic *G. indica*.

## Acknowledgments

We are grateful to the Kerala State Forest Department for granting us permission to conduct the study. We acknowledge funding support from Nature Conservation Foundation, Mysore, CSIR-Centre for Cellular and Molecular Biology, Hyderabad and Science and Engineering Research Board (SERB), New Delhi. We would like to express gratitude to Santhwana and Mahesh for logistical support during the fieldwork in Nelliampathy. We acknowledge the Next-Generation Sequencing (NGS) facility at the CSIR–Centre for Cellular and Molecular Biology (CCMB), Hyderabad, and its staff, especially Subhajit Sen, Tulasi Nagabandi, Mohammad Jafurulla, and Sriram Sudarsanam, for their valuable support in data generation. We also thank Aparna Kumari and Geetha Thanu of the High-Performance Computing (HPC) facility for their assistance with computational resources and support. We are grateful to Shyamal for the illustration that we obtained from Wikimedia Commons, licensed under a Creative Commons Attribution–ShareAlike 3.0 Unported (CC BY-SA 3.0) license.

